# Spatial frequency-based image reconstruction to improve image contrast in multi-offset adaptive optics ophthalmoscopy

**DOI:** 10.1101/2020.12.16.423076

**Authors:** Pedro Mecê, Elena Gofas, Yuhua Rui, Min Zhang, José-Alain Sahel, Ethan A. Rossi

## Abstract

Off-axis detection methods in adaptive optics (AO) ophthalmoscopy can enhance image contrast of translucent retinal structures such as cone inner segments and retinal ganglion cells layer neurons. Here, we propose a 2D optical model showing that the phase contrast produced by these methods depends on the offset orientation. While one axis provides an asymmetric light distribution, hence a high phase contrast, the perpendicular axis provides a symmetric one, thus a substantially lower contrast. We support this model with *in-vivo* human data acquired with a multi-offset AO scanning light ophthalmoscope. Then, using this finding, we provide a post-processing method, named Spatial frequency-based iMAge ReconsTruction (SMART), to optimally combine images from different off-axis detector orientations, significantly increasing the structural cellular contrast of *in-vivo* human retinal neurons such as conne inner segment, putative rods and retinal ganglion cells.

---

Cellular resolution imaging of the living retina can be achieved by combining adaptive optics (AO), which measure and correct for ocular aberrations in real-time [1], with ophthalmoscopes such as confocal scanning-light ophthalmoscope (SLO) [2] and flood-illumination ophthalmoscope (FIO) [3]. AO ophthalmoscopes were first used to detect back-scattered photons to image highly reflective retinal features, such as cones and nerve fiber layer. Later on, by displacing laterally the detection position compared to the illumination, the use of multiply scattered photons was made possible, increasing contrast of translucent retinal structures not previously visualized [4–10]. Different implementations of such off-axis detection methods on both AO-SLO and AO-FIO, as offset aperture, split detection, dark-field, and multi-offset contributed to directly image red blood cells, blood vessel walls, photoreceptor inner segment and ganglion cells *in-vivo*.

Recently, Guevara-Torres *et al*. [11] proposed a one-dimensional geometric optical model for offset aperture in AO-SLO. Their model helps explain the origin of cellular contrast in off-axis detection methods: one edge of the cell appears bright while the opposite edge looks dark. They proposed that when the illumination beam is focused into the edge of a cell, the latter acts as a lens deviating the beam towards or away from an off-axis detector, creating an asymmetric light distribution responsible for the contrast in single cells.

In this Letter, we use the Pittsburgh multi-offset AO-SLO [12], which contains an off-axis detector that can be precisely positioned perpendicularly to the optic axis in eight different positions [Fig. 1(a)], to demonstrate that while one axis will indeed present an asymmetric light distribution generating high cellular contrast, the perpendicular axis has a symmetric light distribution strongly diminishing cellular contrast for this given direction. This behavior is represented in Fig. 1(b) using simple geometric optics and assuming that the cell acts as a weak positive lens (higher refractive index than the surrounding tissue). By exploring this property, we introduce a method to optimally combine images from different aperture positions enhancing structural cellular contrast of human retinal neurons such as photoreceptors and ganglion cells.

**Fig 1.**
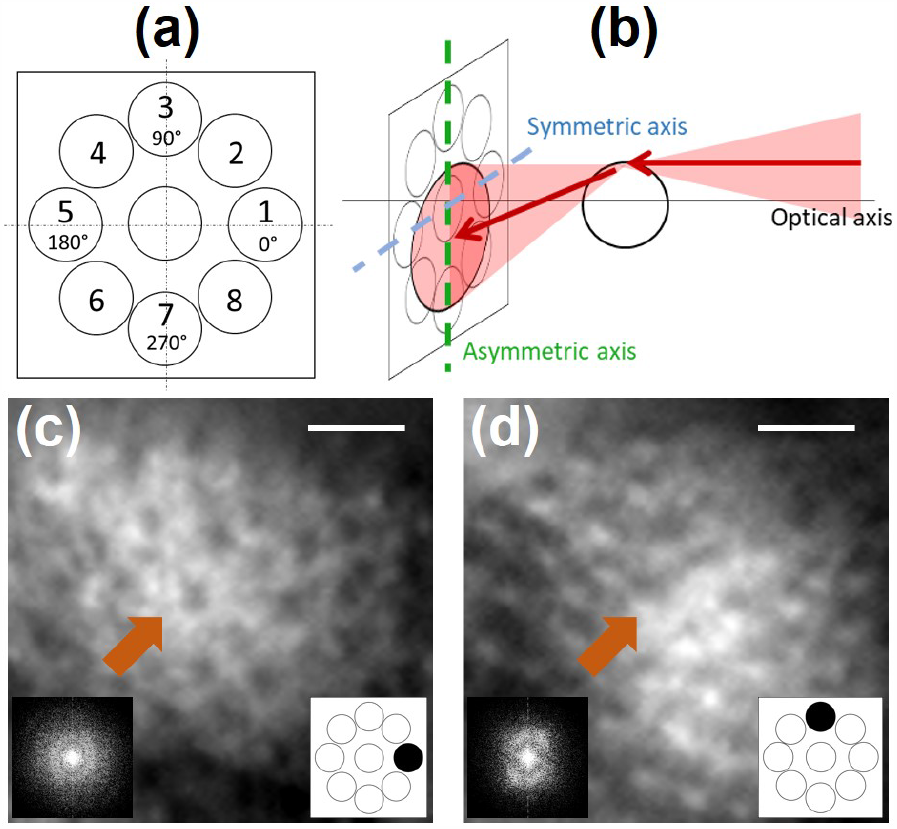
(a) Radial pattern used to sequentially acquire offset aperture images. (b) Geometrical model for multi-offset aperture configuration highlighting the asymmetric and symmetric axis. (c), (d) Magnified images acquired by offset apertures oriented at 0*°*and 90*°*and their respective power spectral density (PSD). Orange arrows point to the same photoreceptor. Note that the cellular contrast and the PSD changes for different offset aperture orientations. Scale bar: 20*µm*.

The Pittsburgh multi-offset AO-SLO was described in details elsewhere [12]. Important for this study, the off-axis detector was placed on motorized xyz stages. After fine alignment and calibration, the detector can be precisely displaced perpendicular to the optic axis in any arbitrary position, enabling acquisition of offset images from different orientations sequentially. For this study we used a radial pattern with displacements of 10 Airy disc diameter (ADD) and a diameter of the detector aperture of 200*µm* (around 7.5 ADD). Retinal images were acquired at 30 Hz in a healthy subject at 10*°*temporal, during 20s for each off-axis detection position. Informed consent was obtained from the subject after the nature and possible outcomes of the study were explained. The experiment was approved by the University of Pittsburgh Institutional Review Board and adhered to the tenets of the Declaration of Helsinki. The subject was seated in front of the system and stabilized with a chin and forehead rest and asked to fixate a target. Eye drops were used for pupil dilation (one drop each of 1% Tropicamide and one 0.5% Phenylephrine). During image acquisition, the total power entering the eye from the AO-SLO and the wavefront sensor beacon were respectively 600*µW* and 20*µW*, which are below the ocular safety limits established by the ANSI. Following acquisition, images were registered using a custom-built strip-based algorithm using the confocal channel image as a reference [13], that was acquired in parallel to the off-axis channel using the same wavelength of light. After registration, image sequences for each offset position were averaged.

To illustrate the symmetric and asymmetric axis, depicted in Fig. 1(b), and their impact on image contrast, we acquired images of the photoreceptor layer using our radially distributed detection pattern [Fig. 1(a)]. Figures 1(c) and 1(d) present two magnified images acquired for offset apertures oriented respectively at 0*°* and 90*°*. Let us consider the case of the 0*°*orientation offset. On the one hand, when illuminating the left and right edges of the photoreceptor, the light beam is deviated respectively towards and away from the given offset aperture, illustrating the asymmetric axis as previously introduced by Guevara-Torres *et al*.. On the other hand, when illuminating the upper and lower edges of the photoreceptor, as the light is deviated in a perpendicular direction (here vertically), this given offset barely sees a change of intensity, hence generating a low contrast and illustrating the symmetric axis. This change of contrast depending on the offset orientation can also be noticed by looking at the power spectral density (PSD) of these images. Indeed, we know that the PSD of photoreceptor mosaic acquired with a back-scattering based AO ophthalmoscope produces the Yellot’s ring [14], *i*.*e*. an isotropically distributed spatial frequency. When computing the PSD for offset apertures oriented at 0*°*and 90*°*, an anisotropic distribution of the photoreceptor mosaic spatial frequency is obtained, and oriented according to the asymmetric axis. **Visualization 1** shows the change of contrast and PSD anisotropy for all eight offset orientations.

To quantify the PSD directionality, we computed the energy around the photoreceptor mosaic spatial frequency, *i*.*e*. the sum of all spatial frequencies between 50 cycles/mm to 200 cycles/mm, as a function of the orientation [Fig. 2(a)]. For a given offset aperture position, the energy is maximal in the asymmetric axis, gradually decreases when the orientation goes away from the asymmetric axis until it reaches a minimum at the symmetric axis. The maximum and minimum points are shifted according to the position of the offset aperture and coincide after a 180*°*rotation (off-axis detector located at opposite position). This behavior can be translated back to the original images, showing that structural cellular contrast is maximal at the asymmetric axis, gradually decreases and reaches a minimal at the symmetric axis. For offset apertures positioned at opposite direction (180*°*from each other), they share the same asymmetric and symmetric axis, but with a negative contrast. Therefore, the difference of opposite offset apertures (henceforth named opposite difference image) optimally enhances structural cellular contrast. However, as previously stated, if only one opposite difference image is considered, the contrast of the cellular structure is only maximal at the asymmetric axis.

**Fig 2.**
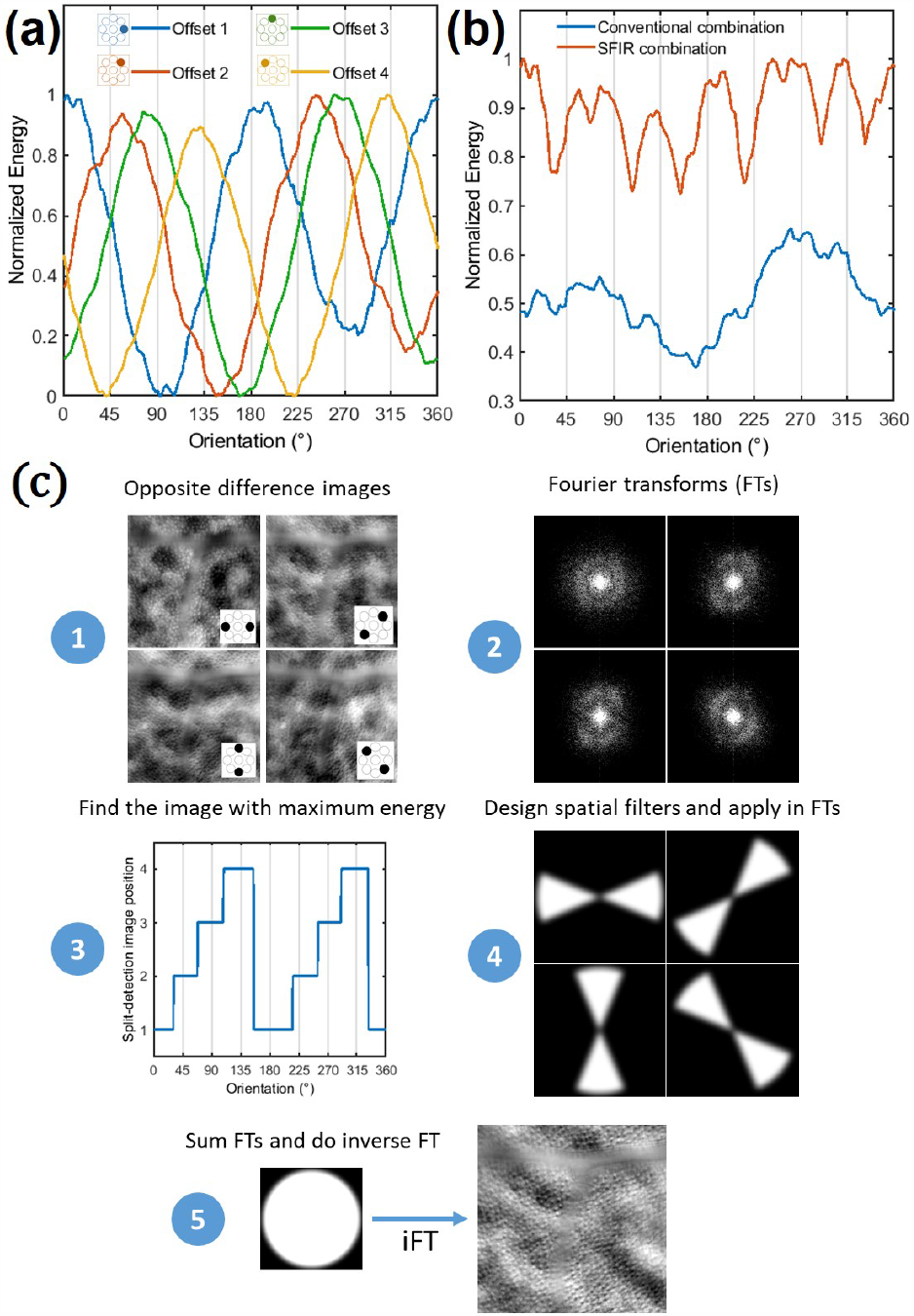
(a) Normalized spatial frequency energy over orientation for offset aperture positions from 1 to 4. (b) Comparison of the normalized spatial frequency energy achieved after a conventional and the SMART combination over orientation, where the expected gain of the SMART method can be appreciated. (c) Post-processing steps of the SMART method.

Conventionally, to mitigate this drawback, previous studies linearly combined opposite difference images from two or several directions [6, 15]. Nevertheless, this approach is not optimal, since it averages the maximum, intermediate and minimum structural cellular contrast for all directions, decreasing the final contrast compared to the maximum reached value at the asymmetric axis. Moreover, since the background spatial frequency information behaves independently of the offset aperture orientation, a linear combination of opposite difference images will favor background signal (low spatial frequency), drastically decreasing the signal-to-background ratio of the final image. Instead of linearly averaging opposite difference images, we propose to only use spatial frequencies close to the asymmetric axis, where the contrast is maximum, which corresponds in obtaining the superior envelope of Fig. 2(a). We call this proposed approach Spatial frequency-based iMAge ReconsTruction (SMART) method. Figure 2(b) presents a comparison of the final spatial frequency energy obtained over orientation using the proposed SMART combination method and the conventional one. The SMART method steps can be decomposed in five steps: (1) Generate opposite difference images; (2) compute their Fourier Transform (FT); (3) for a range of spatial frequency of interest, compute the energy over the orientation, and find the opposite difference image which carries the highest energy for a given orientation; (4) based on the previous information, create a mask (spatial filter) for each opposite difference image FTs; (5) after applying a Gaussian filter in each mask (to avoid artifacts), apply the corresponding spatial filters in each of offset image FTs, reconstruct the final image by summing the opposite difference image FTs, and do the inverse FT. Figure 2(c) presents a schematic of the required steps to apply the SMART method.

Figures 3(a) and 3(b) show the enhancement of image contrast of the photoreceptor layer when applying the SMART method in comparison to the conventional image combination. Plots of image intensity along drawn lines 1, 2, 3 and 4 are shown in Fig. 3(c). In all presented cases, a notable contrast gain is observed. The use of the SMART method enhanced the Michelson contrast of drawn lines 1, 2, 3 and 4 by a factor of 1.87, 1.65, 1.77 and 1.8 respectively. Figure 3(d) presents the PSD for the conventional and the SMART combination cases. Whilst the low-spatial frequency power is kept unchanged (background content), the photoreceptor mosaic spatial frequency content (highlighted by the pale green column) presents an enhancement of a factor of 2.63. Owing to the contrast improvement through the SMART method, putative rods become visible, as it is highlighted by red arrows in Fig. 3(c) (see **Visualization 2** for magnified images).

**Fig 3.**
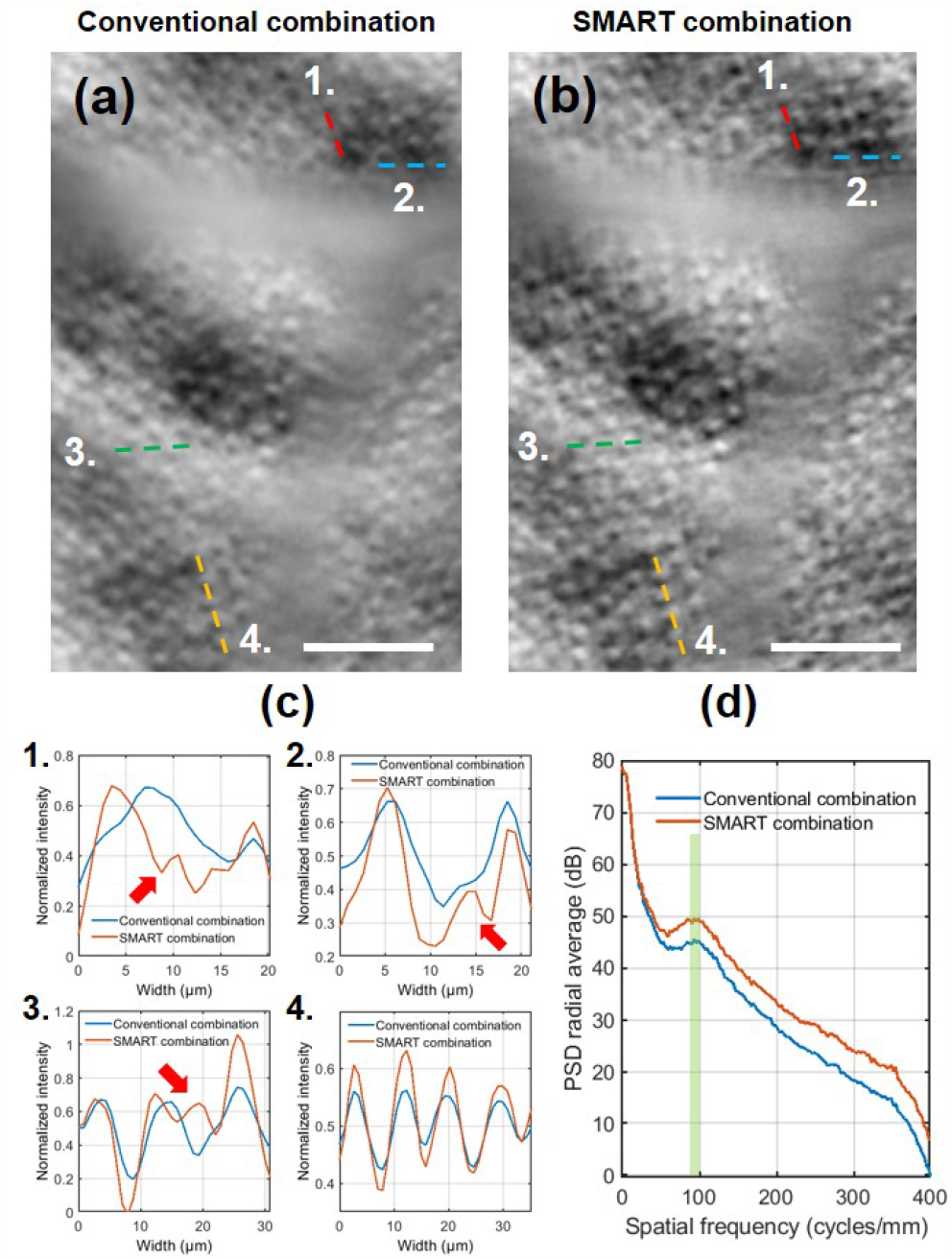
Multi-offset AO-SLO photoreceptor layer image after (a) conventional and (b) SMART combination. (c) Plots of drawn lines in (a) and (b) highlighting the contrast gain with the SMART method. Note in plots 1., 2. and 3. that owing to the SMART method putative rods, invisible when using the conventional combination, become visible (red arrows). See **Visualization 2** for magnified images where putative rods become visible. (d) Computed radial average PSD after conventional and SMART combination, where the pale green column highlights the contrast improvement in the range of photoreceptor mosaic spatial frequency. Scale bar: 50*µm*.

Enhancing phase contrast in off-axis detection methods becomes crucial when imaging weakly scattering structures such as *in-vivo* human retinal ganglion cells. Although we recently demonstrated an improved image contrast of *in-vivo* human retinal ganglion cells using a multi-offset aperture AO-SLO 4(a) [12] compared to the work of Rossi *et al*. [6], there was still regions where we could not clearly identify the cells because of low contrast.

Using the completely digital SMART method, we were able to increase the image contrast, as it is highlighted in Fig. 4 and **Visualization 3**. Magnified regions 1, 2, 3 and 4 and plots of image intensity along drawn lines are shown in Fig. 4(d). In these four cases, the Michelson contrast improved by a factor of 2.3, 1.95, 1.5 and 1.42 respectively. Figure 4(c) presents the PSD for the conventional and the SMART combination. Once again, one can notice that while low spatial frequency coming from the background is unchanged, high spatial frequencies carrying the structural cellular contrast of ganglion cells (pale green column) are enhanced by a factor of 2.88.

**Fig 4.**
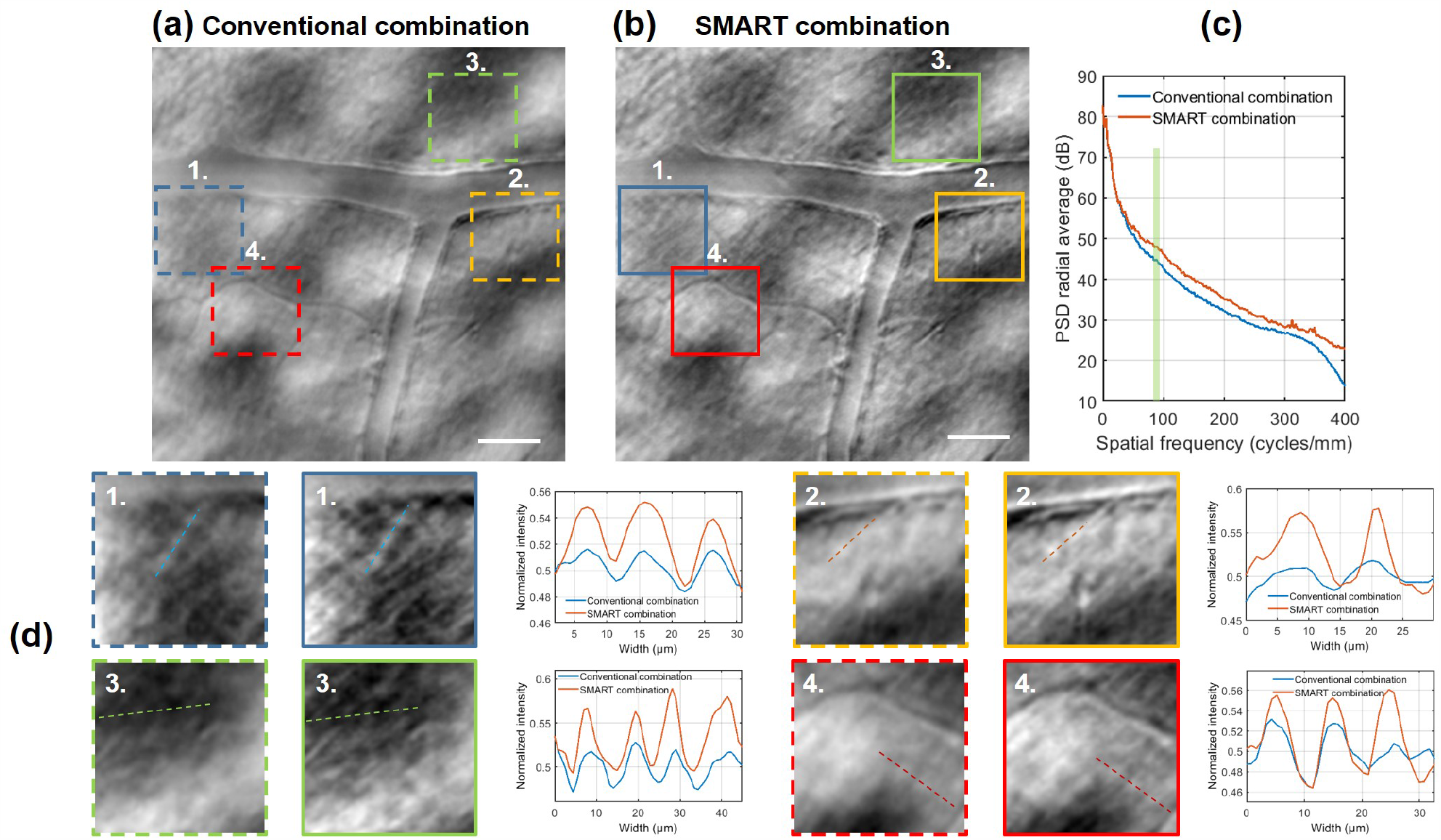
Multi-offset AO-SLO ganglion cell layer image after (a) conventional and (b) SMART combination. (c) Computed radial average PSD after conventional and SMART combination, where the pale green column highlights the contrast improvement in the range of ganglion cells mosaic spatial frequency. (d) Magnified regions of 70*µm×*70*µm* and plots of drawn lines highlight the contrast enhancement due to the SMART method. The gain in terms of contrast is also visible in **Visualization 3**. Scale bar: 50*µm*.

The proposed two-dimensional (2D) optical model, taking into account an asymmetric and symmetric axis, carrying the maximum and the minimum phase contrast, helps to explain the fact that blood vessels seem to disappear according to the offset aperture orientation [6, 8]. **Visualization 4** presents an example on the influence of offset aperture orientation on blood vessel wall contrast. Considering the case of the horizontal blood vessel, the light beam is deviated by the blood vessel wall in a vertical direction, provoking an asymmetric light distribution, and maximizing the contrast in off-axis detectors positioned in this axis. Contrarily, off-axis detector positioned in the symmetric axis will barely see a change of intensity, and no contrast would be produced for this vessel. Through the use of the SMART method, all capillaries, independently on their orientation, present an optimal contrast [see Fig. 4(b)], which could be extremely helpful when measuring the wall-to-lumen ratio and other important vessel-based biomarkers [16]. Although not shown here, the SMART method might also improve contrast of generated perfusion maps using off-axis detection methods [4].

In this Letter, we proposed a 2D model, complementary to the 1D model presented by Guevara-Torres *et al*. [11], showing that besides the existence of an asymmetric axis, where the phase contrast is maximal, when moving away from this axis, the structural phase contrast diminishes, until reaching a minimum contrast in a symmetric axis. Based on this behavior, we proposed the SMART method, which optimally combine offset aperture images acquired in different orientations, by smartly selecting spatial frequencies presenting maximal energy, thus enhancing the structural cellular contrast of photoreceptor inner segment, putative rods, and ganglion cells. Still, there is some room to improve phase contrast using off-axis detector techniques, especially during acquisition. Although not shown yet in humans, conjugating the off-axis detector to a deeper layer [11] seems to be a promising strategy to maximize the phase contrast in the asymmetric axis. Combining this strategy with a higher AO-loop rate [17], to correct for dynamic ocular aberrations, and the SMART method would enable a maximal contrast in all directions. Additionally, finely probing different offset orientations (here we were limited to eight positions) should improve even more the phase contrast through the SMART method.

## Acknowledgments

The authors want to thank Kate Grieve and Shan Suthaharan for fruitful discussion.

